# *Salmonella* relies on siderophore exploitation at low pH

**DOI:** 10.1101/2025.05.06.652355

**Authors:** Manon Ferry, Connor Sharp, Isabelle J. Schalk, Olivier Cunrath

## Abstract

*Salmonella enterica*, a prominent enteric pathogen, employs sophisticated iron acquisition mechanisms to overcome host-imposed iron limitation, notably through the production and uptake of siderophores—small, high-affinity iron-chelating compounds that scavenge iron from the host environment. In this study, we investigate how environmental pH influences *Salmonella’s* preference for its endogenous siderophores versus exogenous siderophores within the physiological range of the gastrointestinal tract. Through competition assays, gene expression analysis, and siderophore quantification, we demonstrate that *Salmonella* increasingly relies on exogenous siderophores under acidic conditions. This shift is attributed to reduced production of its endogenous siderophores, enterobactin and salmochelin. Deletion of the sigma factor RpoS enhances endogenous siderophore production and iron acquisition at low pH, suggesting its role in regulating iron homeostasis. Our findings reveal a pH-dependent adaptation in *Salmonella’s* iron acquisition strategy, highlighting the pathogen’s versatility in nutrient acquisition across varying gastrointestinal conditions. This research provides insights into *Salmonella’s* pathogenicity and may inform the development of targeted interventions for *Salmonella* infections.

## Introduction

Iron is a vital nutrient for all organisms, playing a key role as a cofactor in enzymes that drive growth and metabolism^1^. However, iron bioavailability is limited in many environments, making it a scarce resource for both the host and invading pathogens^2–4^. During infection, the host organism mounts a defense strategy known as nutritional immunity, which tightly restricts the levels of available iron, effectively limiting its accessibility to pathogens^5,6^. By limiting access to iron, the host forces pathogens to compete for this essential resource. This competition shapes the course of infection and drives pathogens to develop various strategies for acquiring the iron they need to survive and proliferate. One of these strategies is the production of siderophores—high-affinity molecules that scavenge iron from the environment and facilitate its uptake into bacterial cells^7^. Siderophores are synthesized by various microorganisms in response to iron scarcity and are secreted into the extracellular environment, where they form stable complexes with iron^8^. Once the siderophore-iron complex is formed, it is recognized and actively transported back into the microbial cell via specific transport systems. These transporters, located on the microbial surface, are highly selective and efficiently bring the iron-bound siderophore inside the cell, where the iron is released for cellular use^9^.

Numerous studies highlight the essential role of siderophores in the inflamed intestinal environment^10–13^. The host actively interferes with bacterial siderophore utilization by producing molecules like lactoferrin, transferrin and calprotectin, which tightly bind iron, limiting its availability to pathogens^14–16^. Additionally, the secretion of lipocalin-2 further restrict microbial access to iron, by capturing some iron-loaded siderophores^17^. Beyond the host’s defense mechanisms, the gut microbiota plays an additional role in the competition for iron. Certain commensal bacteria, such as *Lactobacillus* and *Bifidobacterium* species, along with some fungal strains like *Aspergillus* and *Malassezia*, are known to produce their own siderophores to acquire iron^18–21^. Siderophores can also enter the intestinal environment through the diet, as they are produced by microorganisms naturally found in fermented foods^21,22^. Together, these factors suggest that the intestinal environment is a dynamic landscape where iron is bound to a variety of siderophores. To thrive in this complex setting, many bacteria have evolved not only to produce their own siderophores but also to exploit exogenous siderophores — siderophores produced by other organisms and present in their environment (also referred to as xenosiderophores) ^23–25^. For instance, commensal gut bacteria such as *Bacteroides* can exploit siderophores produced by other microorganisms, as can gut pathogens like *Candida albicans*, *Clostridioides difficile*, and *Escherichia coli*^26–29^. This dual capability to both produce and exploit siderophores provides pathogens with a versatile strategy to navigate the diverse ecological niches of the gut. To explore how these strategies may influence pathogen proliferation, we focus on *Salmonella enterica enterica* serovar Typhimurium, a prominent model pathogen of gastrointestinal infections and iron homeostasis^13^.

*Salmonella* is capable of producing two catecholate-type siderophores—enterobactin (ENT), through the biosynthetic gene cluster EntCEBAH^30,31^, and salmochelin (SLC), its C-glucosylated derivative, through glycosylation mediated by the IroB enzyme^32^—in response to iron scarcity^33^. Of all reported siderophores so far, ENT exhibits the highest affinity for iron (K*_a_* = 10^49^ M^-1^)^34^ but can be neutralized by the host protein lipocalin-2, which sequesters ENT, thus blocking this iron acquisition pathway^17^. In contrast, the glycosylation of SLC (K*_a_* = 10^45^ M^-1^)^12^ prevents recognition by lipocalin-2, allowing *Salmonella* to evade this immune defense while maintaining its high affinity iron acquisition system, even under conditions of severe iron limitation^35^.

As for all Gram negative bacteria, ferri-siderophore complexes formed in the bacterial environment cross the outer membrane through TonB-dependent transporters (TBDTs), utilizing energy derived from the proton motive force transmitted by the TonB-ExbB-ExbD inner membrane complex^36–40^. TBDTs are highly specific to their ligand, ensuring selective uptake of the iron-loaded siderophores (recently reviewed here^8,13^). In *Salmonella*, ferric-ENT, ferric-SLC, and their ferric break-down products 2,3-dihydroxybenzoylserine (DHBS) are recognized by specific the following TBDTs: FepA for ENT and DHBS, IroN for SLC and DHBS, and CirA for DHBS (Fig. 1, left part)^31,41,42^. Beyond producing its own endogenous siderophores, *Salmonella* can also exploit three exogenous siderophores found in its environment: ferrichrome (FC) (K*_a_* = 10^29^ M^-1^)^23,43^, coprogen (CPG) (K*_a_* = 10^27^ M^-1^)^36,44^, and ferrioxamines (DFO) (K*_a_* = 10^32^ M^-1^)^37,45^, all of which are hydroxamate-type siderophores (Figure 1, right part). This specialized transport mechanism enables *Salmonella* to efficiently acquire iron from the environment, particularly under iron-limited conditions, by facilitating the uptake of the strong affinity ferric-siderophore complexes.

Genes encoding siderophore transporters or involved in their biosynthesis are negatively regulated by the ferric uptake regulator Fur, a globally conserved transcriptional regulator that controls iron homeostasis in bacteria^46^. Fur functions as a repressor by binding to specific DNA sequences known as Fur boxes located in the promoter regions of target genes when complexed with Fe²⁺. In addition to the regulation by Fur, certain genes involved in siderophore acquisition are controlled by other mechanisms. For example, the TBDT FoxA, responsible for ferrioxamine E (DFOE) uptake, exhibits a ligand-specific upregulation mechanism mediated by the AraC-type transcriptional regulator FoxR. FoxR binds to DFOE in the cytoplasm, activating the transcription of the *foxA* gene and thereby upregulating FoxA expression to improve *Salmonella*’s access to this exogenous siderophore (Fig. 1, right part), thus optimizing the bacterial iron acquisition process under iron-limited conditions^47^.

Our understanding of how pathogens selectively use siderophores under various infection conditions remains limited. One key question is whether pathogens primarily rely on the production of their own siderophores or whether they exploit exogenous ones, and how this choice is influenced by the dynamic conditions of the intestinal environment. Environmental factors such as changes in pH, which fluctuate within the gut, could significantly impact iron availability and may affect the way siderophores are utilized^39,48–50^. Given the energetic cost of siderophore production and the nutrient restrictive environment imposed by both the host and the innate microbiome, it is plausible to assume that, under certain conditions, pathogens might favor the exploitation of exogenous siderophores^51^. This raises important questions about the adaptive mechanisms’ pathogens employ to optimize their iron acquisition in the face of a constantly shifting gut environment. Here we show that siderophore exploitation is a vastly conserved strategy among all tested *Salmonella* sub-populations. We further show that exogenous siderophore exploitation is key for *Salmonella’s* survival during iron starvation at low pH. Despite complex regulatory mechanisms involved in iron homeostasis, we find that this pattern can be explained by a simple underlying principle: a stark reduction of the endogenous siderophore production in acidic conditions. Furthermore, we have shown that this down-regulation may represent a pathogen’s weakness that can be utilized for better pathogen control strategies^21,52–54^.

**Figure 1.**
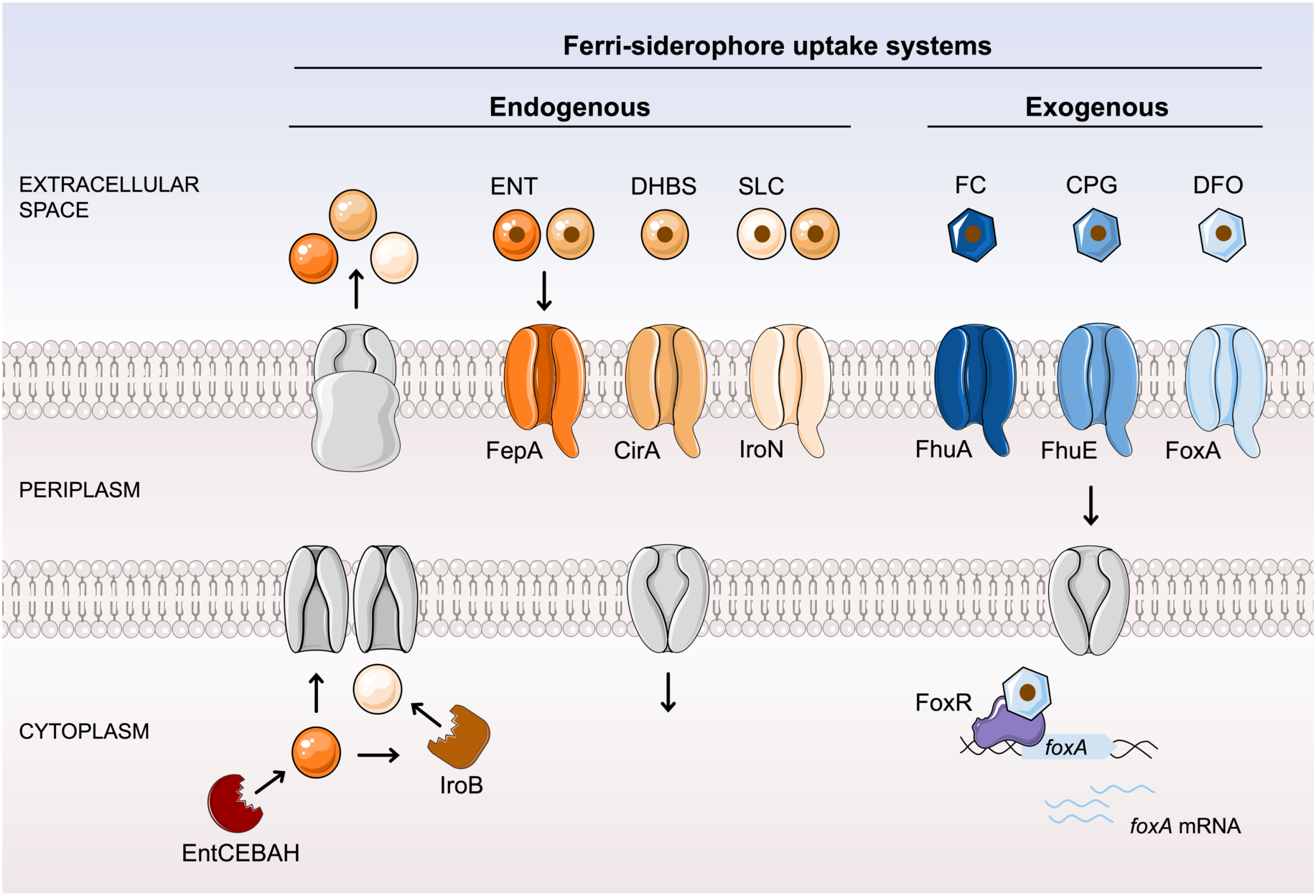
Siderophore mediated iron uptake in *Salmonella enterica*. Schematic diagram illustrating the pathways for the acquisition of endogenous (left) and exogenous (right) siderophores identified in *S. enterica* Typhimurium. Iron (brown circle) is captured from the extracellular environment and the ferri-siderophore complex is transported into the cell by TBDTs. Endogenous siderophores, produced by the *entCEBAH* operon, include ENT (dark orange circle), its break-down product DHBS (medium orange circle), and SLC (light orange circle). SLC is a glycosylated derivative of ENT, with glycosylation facilitated by IroB. All three are recognized by TBDTs FepA, IroN, and CirA. Exogenous siderophores—FC (dark blue hexagon), CPG (medium blue hexagon), and DFO (light blue hexagon)—are imported by TBDTs FhuA, FhuE, and FoxA, with FoxA being positively regulated by the cytoplasmic regulator FoxR.

## Results

### Siderophore uptake systems are very conserved suggesting key ecological roles in *Salmonella enterica* lifestyle

*Salmonella enterica* is well known for its ability to both produce its own siderophore and exploit exogenous siderophores. *Salmonella* is able to synthesize ENT through the biosynthetic gene cluster EntCEBAH^30,31^ and SLC through a glycosylation of ENT mediated by the IroB enzyme^32^. ENT, SLC, and their break down product DHBS are recognized by specific TBDTs: FepA for ENT and DHBS, IroN for SLC and DHBS, and CirA for DHBS^31,41,42^. To assess the conservation of *Salmonella’s* endogenous siderophore uptake systems we analyzed siderophore-related genes in 183 *S. enterica* strains. These strains were randomly chosen with one strain selected for each unique multilocus sequence typing (MLST) profile on the PATRIC database using Python’s *random.choice()* function to ensure a diverse dataset^55^. Our results reveal a strong conservation of all endogenous siderophore biosynthesis and uptake systems across the species (Fig. 2). Nearly all strains possess the *entC* and *fepA* genes (>99%), which enable the biosynthesis and uptake of ENT, respectively, highlighting the importance of these genes in *Salmonella*’s iron acquisition strategies. Likewise, *cirA* is universally present, supporting the consistent use of this transporter across strains. Furthermore, the high conservation of both *iroB* and *iroN* (98.9%) in almost all strains suggest a strong conservation of the capacity to produce and import SLC.

In addition to its own siderophores, *Salmonella* can exploit exogenous siderophores present in its environment, such as FC, CPG, and DFO. These are imported via other TBDTs: FhuA for FC, FhuE for CPG, and FoxA for DFO^36,37,39^. Recent research has further demonstrated the regulatory role of the AraC-type regulator FoxR in enhancing DFOE uptake^47^. When analyzing the conservation of genes encoding for exogenous siderophore TBDTs as described above, we could show that although their conservation was slightly more variable than for the endogenous transporters, conservation still remains high (Fig. 2). The *fhuA* gene, responsible for FC transport, is slightly less conserved but is still present in about 73% of the strains. Conversely, the genes for CPG and DFO transport—*fhuE* (> 99%) and *foxA* (94%), respectively—are highly conserved, as is *foxR* (95%), the positive regulator of *foxA*. All but one strain possesses at least one exogenous siderophore transporter, underscoring the species-wide dependency to access exogenous siderophores in diverse environments (Fig. 2). A broader analysis of approximately 1,200 strains with unique genomes showed the same trend, reinforcing the conclusion that siderophore-dependent iron uptake systems are highly conserved across the species and likely play a key role in *Salmonella*’s infection process (fig. S1; Tab S5).

**Figure 2.**
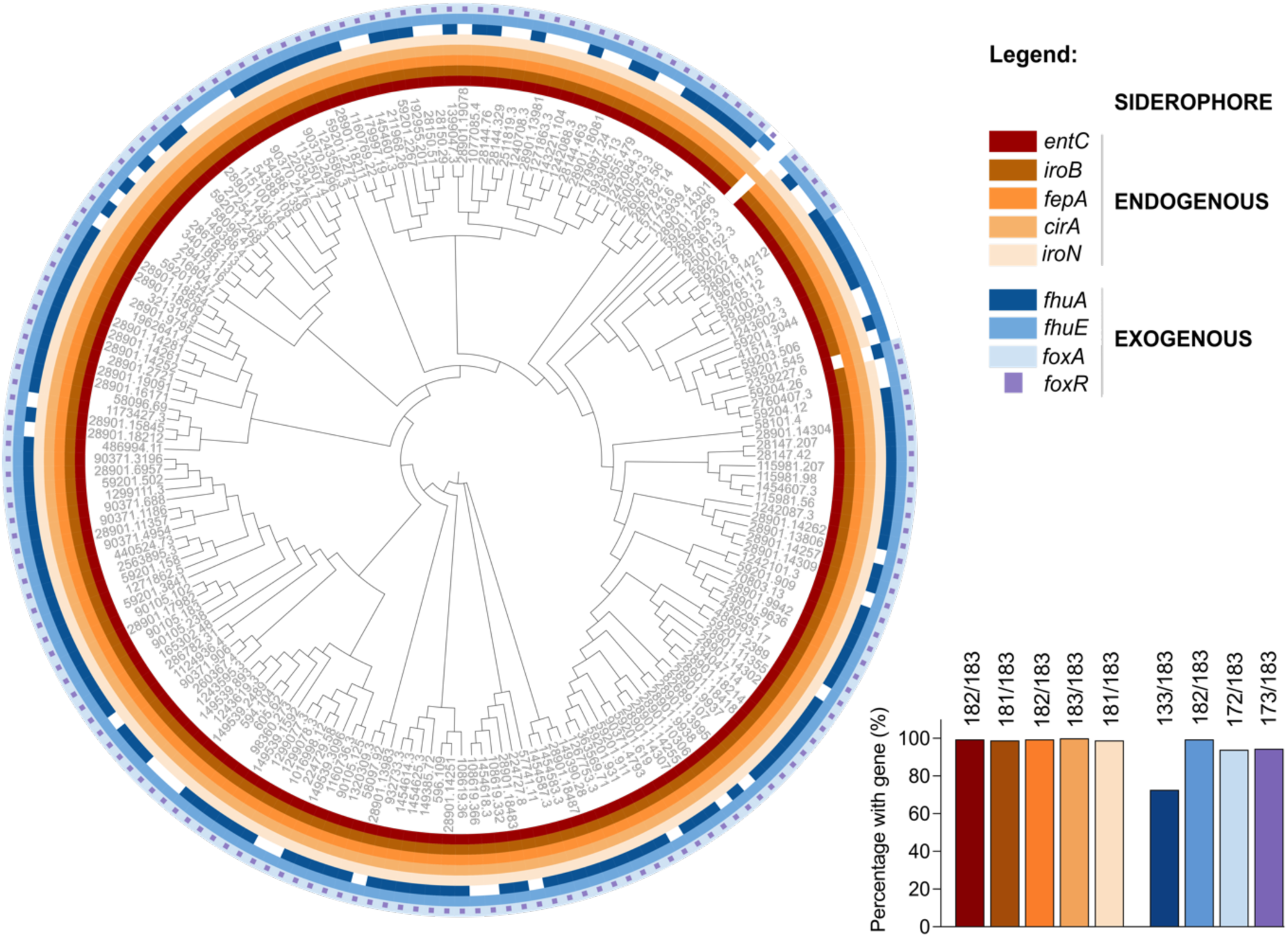
Conservation of siderophore-related genes across *Salmonella enterica* species highlights their ecological importance. Phylogenetic tree of 183 *Salmonella* strains. The phylogeny is based on the core gene alignment of all strains, selected to maximize genetic diversity with each having a unique MLST profile. Circles around the tree depict the presence or the absence of specific genes, with each circle corresponding to one gene. Colored rectangles next to each strain’s Genome ID (PATRIC BV-database) indicate gene presence, while white rectangles show absence. The histogram shows the percentage of strains that possess each studied gene.

### The reliance of *Salmonella* on exogenous siderophore exploitation increases in acidic conditions

Changes in pH within the gastrointestinal tract can significantly affect siderophore physicochemical properties^48,56^. As the pH fluctuates, the chelating properties of siderophores may be modified, which in turn may influence their relative importance in iron acquisition. For example, ENT, a catecholate-type siderophore, sees its affinity strongly decrease upon acidification of the environment, due to protonation of the catechol groups ^34,48^. On the other hand, hydroxamate siderophores like deferoxamine (DFO) depict a weaker affinity for iron at neutral pH but show a weaker decrease in affinity upon acidification^48,57^. Given this pH-dependent variability in siderophore affinity, we hypothesize that acidic conditions in the gut could influence *Salmonella*’s preference towards exploiting exogenous hydroxamate siderophores, as their uptake might be less affected by the changes in iron affinity compared to their endogenous siderophore ENT.

To investigate the influence of pH on *Salmonella*’s use of exogenous siderophores, we performed bacterial competition assays (Fig. 3A). In these assays we investigated two different exogenous siderophores uptake systems; the FC uptake, in which the transcription and consequently the expression of the TBDT FhuA is solely regulated by the general iron regulator Fur, and the iron uptake by DFOE, where the transporter FoxA has its expression regulated by Fur and is also inducible by the presence of the DFOE via FoxR (Fig. 1)^37,47^. A WT strain and an isogenic strain lacking the gene encoding for the exogenous-siderophore transporter (*fhuA* or *foxA*) or the regulator (*foxR*) were co-cultured in iron-depleted media at various pH levels, with increasing concentrations of the corresponding exogenous siderophores. It is important to note that both competing strains were capable of producing and using their endogenous siderophores to acquire iron. For each condition, we measured the competitive index (CI) of the mutant strain relative to the WT strain (as calculated by the CFU ratio (mutant/WT) at 48h divided by the CFU ratio (mutant/WT) of the inoculum) to assess the advantage conferred by the ability to exploit exogenous siderophores across different pH levels (Fig. 3A). These experiments showed a decreased fitness of the mutant strain in increasing concentrations of our exogenously added siderophores (Fig. 3BCD). At intermediate concentration such as 0.1 μM this decrease was significantly more pronounced at low pH. Therefore, our results showed a significant pH-dependent effect on the advantage provided by FC exploitation (Fig. 3B). The inability to use FC, due to the deletion of *fhuA*, demonstrated that *Salmonella* gains a distinct advantage from FC exploitation more quickly in acidic conditions than at neutral pH. Similarly, when comparing the WT strain that can use DFOE to the *foxA* mutant strain that cannot, we observed the same pH-dependent advantage (Fig. 3C). Since FoxA appears more important under acidic conditions, we investigated the relative advantage of FoxR-mediated upregulation of the FoxA transporter. Our findings indicate that, in the presence of all tested concentrations of DFOE, FoxR-mediated upregulation is only relevant for *Salmonella* in acidic conditions (Fig. 3D). Growth assays revealed a striking pH-dependent role for FoxR. At neutral pH, a *foxR* mutant grows similarly to WT, even in the presence of 10 µM of DFOE, indicating that FoxR is dispensable under these conditions (fig. S2). In contrast, at acidic pH, the same *foxR* mutant exhibits a significant growth defect compared to WT when DFOE is present, demonstrating that FoxR is essential for *Salmonella’s* fitness in acidic environments (fig. S2). The same findings also hold in an *entC* mutant background, unable to produce its own siderophore (fig. S3). These findings reveal that acidic pH represents a condition in which the regulator FoxR becomes crucial for iron acquisition. At low pH, *Salmonella* requires both the FoxA transporter and its up-regulation by FoxR to effectively exploit DFOE siderophore; without which siderophore exploitation may be blocked.

**Figure 3.**
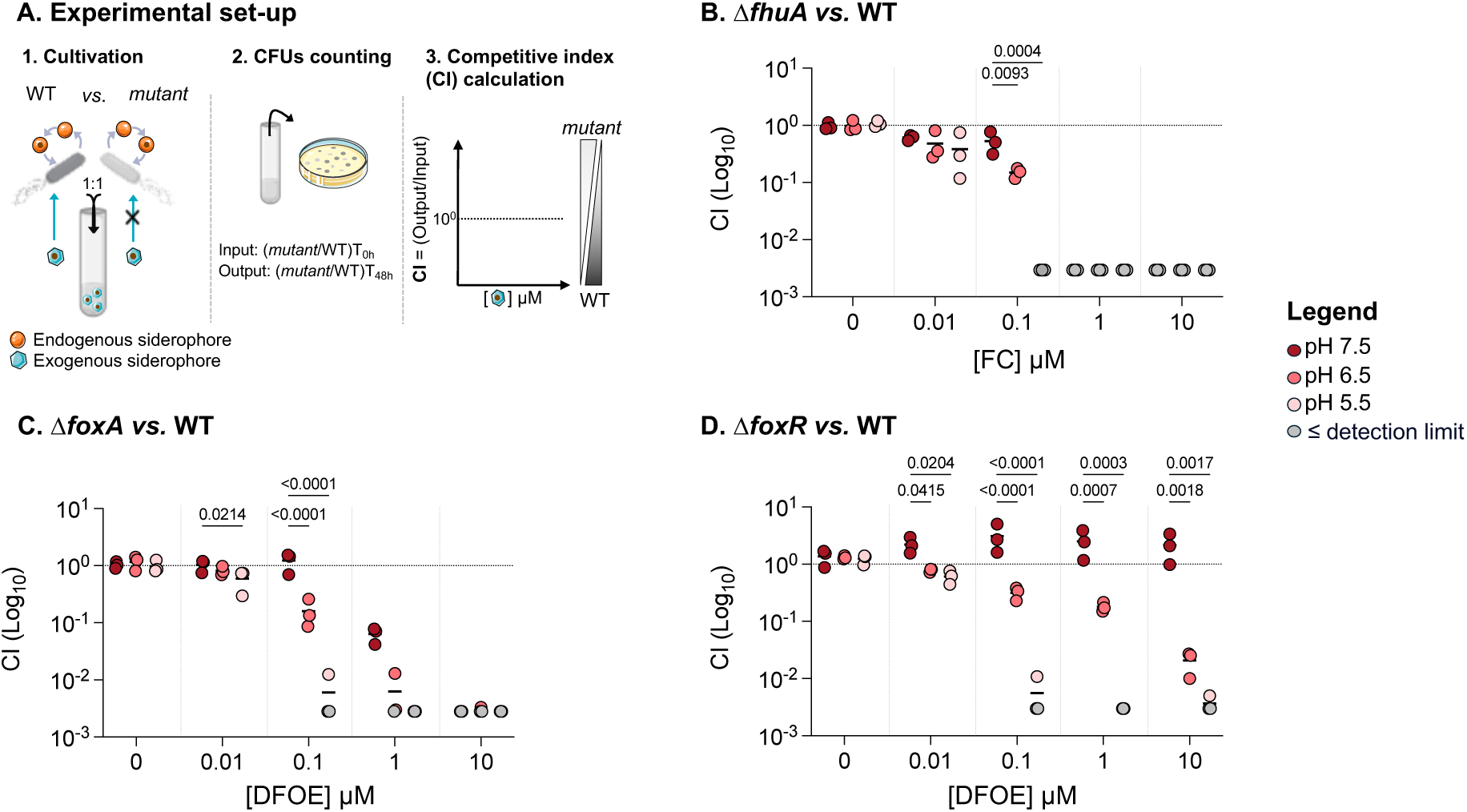
The reliance of *Salmonella* on exogenous siderophore exploitation increases in acidic conditions. **A**. Experimental setup for competition assays. Two fluorescently labeled *Salmonella* strains are mixed in equal amounts, with one strain unable to utilize the exogenous siderophore. Cultures are grown in Minimal Medium (MM) with or without the exogenous siderophore. After 48 hours, the ratio of mutant to wild-type colonies is determined by plating on selective agar, and the competitiveness index (CI) is calculated (for more details the Method section). A CI > 1 indicates a competitive advantage for the mutant strain, while a CI < 1 indicates an advantage for the wild-type strain. **B.** Competitive indexes of Δ*fhuA* s *vs*. WT strain in presence of FC. **C.** Competitive indexes Δ*foxA vs*. WT strain in presence of DFOE. **D.** Competitive indexes of Δ*foxR vs*. WT strain –regulator of FoxA in presence of DFOE. All assays were performed in MM at pH 7.5 (dark red), 6.5 (medium red), or 5.5 (light red), with or without increasing concentrations of exogenous siderophores. The output ratio was calculated after 48 hours of culture. Each data point represents one biological replicate (n = 3). Points below the detection limit for each assay are shown in gray. Statistical analysis was performed using one-way ANOVA followed by Tukey’s multiple comparisons test. Exact p-values are reported for all comparisons with p < 0.05 considered statistically significant.

### Reduced endogenous siderophore production increases dependence on exogenous siderophores in acidic conditions

While the affinity between iron and the siderophore varies at different pH, we explored if our pH-dependent phenotype could be explained by more than just the affinity difference between iron and the siderophores^48^. TBDTs that import siderophores are highly specific, with each transporter recognizing a particular siderophore-iron complex^8^. Previous studies have shown that pH can alter the affinity of the siderophore-iron complex for its transporter^51^. To understand why *Salmonella* gains a significant growth advantage from exogenous siderophore utilization in acidic conditions, we examined whether pH affects the affinity of the siderophore-iron complex for its transporter. Therefore, we measured the affinity of Fe-DFOE, one of the three exogenous siderophores utilized by *Salmonella*, for its transporter FoxA. Specifically, *Salmonella* cells expressing FoxA were incubated with increasing concentrations of radiolabeled ^55^Fe-DFOE at 4°C. At this temperature, Fe-siderophores bind to their TBDT but no uptake into the bacteria occurs^58^. After incubation, unbound complexes were removed by filtration and the radioactivity associated with the bacterial pellet was measured by scintillation counting. These data were used to generate binding curve and calculate the dissociation constant (K*d*), providing a quantitative measure of the interaction between Fe-DFOE and FoxA. Our findings indicate that the growth advantage is not due to changes in siderophore binding properties, as DFOE displayed consistent affinity for its transporter FoxA at both pH 7.5 and 5.5 with dissociation constants of around 3.2 ± 7.6 nM and 3.2 ± 11.0 nM, respectively (fig. S4).

Another possibility is that the increased dependence on exogenous siderophore under acidic condition could be attributed to modifications in bacterial physiology induced by pH. The expression profile of the *ryhB2* reporter —a small regulatory RNA involved in iron homeostasis— across our pH range (5.5, 6.5, 7.5) indicates that the bacterial iron requirement is reduced in acidic environments. This may partly explain the downregulation of siderophore systems observed under these conditions, while also confirming that all tested conditions remain iron-deleted (fig. S5). We then explored two possible scenarios to explain the increased reliance on exogenous siderophores: either exogenous uptake systems are more highly expressed in acidic environments, or endogenous uptake systems are less expressed, leading to greater dependence on exogenous siderophore exploitation. To assess the expression levels of siderophore-uptake pathways in relation to pH, we used plasmid-encoded fluorescent transcriptional reporter fusions^59,60^ in which we have cloned the promoter region of the gene of interest in front of GFP.

Using this approach, we observed a reduction in the transcription of the exogenous *foxA* and *fhuA* genes under acidic conditions, even when grown in the presence of their respective ligand (Fig. 4A). We then investigated the expression of genes involved in the endogenous siderophore uptake system (Fig. 4B). Specifically, the expression of the *entCEBAH* operon, which is responsible for ENT production, and *iroB*, which contributes to SLC production, was also reduced in acidic conditions. Similarly, the expression of endogenous siderophore TBDTs such as *cirA*, *fepA*, and *iroN* decreased, suggesting that endogenous siderophores are produced at lower levels in acidic conditions. This was further supported by the quantification of endogenous siderophores using UV-visible spectroscopy, which revealed reduced concentrations of endogenous siderophores (ENT/SLC and DHBS) in the supernatant of strains cultivated under acidic conditions (Fig. 4C). To make sure that this phenomenon is not strain-specific, we quantified ENT/SLC and DHBS production in various phylogenetically diverse *Salmonella* strains, which reveal similar trends (fig. S6). In addition to the use of siderophores, *Salmonella* also possesses ferrous iron uptake systems such as FeoABC, SitABCD and MntH ^61–63^ (recently reviewed here^13^). To investigate if those system may be of greater importance in acidic conditions, we also examined whether the expression of the three major ferrous iron import systems are upregulated under acidic conditions. However, our results indicate that these import systems are not significantly more expressed in acidic environments compared to neutral ones (fig. S7). Taken together, these findings suggest that, under acidic conditions, *Salmonella* shifts toward a greater reliance on exogenous siderophores due to reduced production of its own endogenous siderophores and a change in iron-chelating affinity.

At neutral pH, ENT/SLC siderophores have a higher affinity for iron than hydroxamate siderophores, but at acidic pH, their affinity decreases to a level comparable to that of hydroxamates, making them less competitive in acidic conditions^48^. In our competition model, hydroxamate siderophores are already present in the environment, which may further enhance *Salmonella*’s dependence on these exogenous siderophores to access iron. This combination of limited production and reduced affinity in acidic conditions likely drives *Salmonella* to favor exogenous siderophore uptake. To test whether this increased reliance is indeed due to reduced endogenous siderophore production, we supplemented the culture medium with pure ENT to see if this addition could alleviate the dependence on exogenous sources (Fig. 4D). We competed a WT strain with a strain deficient in DFOE uptake (*foxA* mutant) at both pH 7.5 and 5.5. While the addition of DFOE alone conferred a growth advantage to the WT strain in acidic conditions, this advantage was lost when the same concentration of ENT was added alongside DFOE. The presence of exogenous ENT effectively reduces *Salmonella*’s reliance on exogenous siderophores, highlighting the critical role of endogenous siderophore production in facilitating growth under acidic conditions.

Our results indicate that *Salmonella*’s increased reliance on exogenous hydroxamate-type siderophore exploitation under acidic conditions is driven by a combination of factors. The decreased production of endogenous catechol siderophores, combined with the reduced affinity of *Salmonella*’s endogenous siderophores for iron in acidic environments, drives a greater dependence on exogenous sources of siderophores.

**Figure 4.**
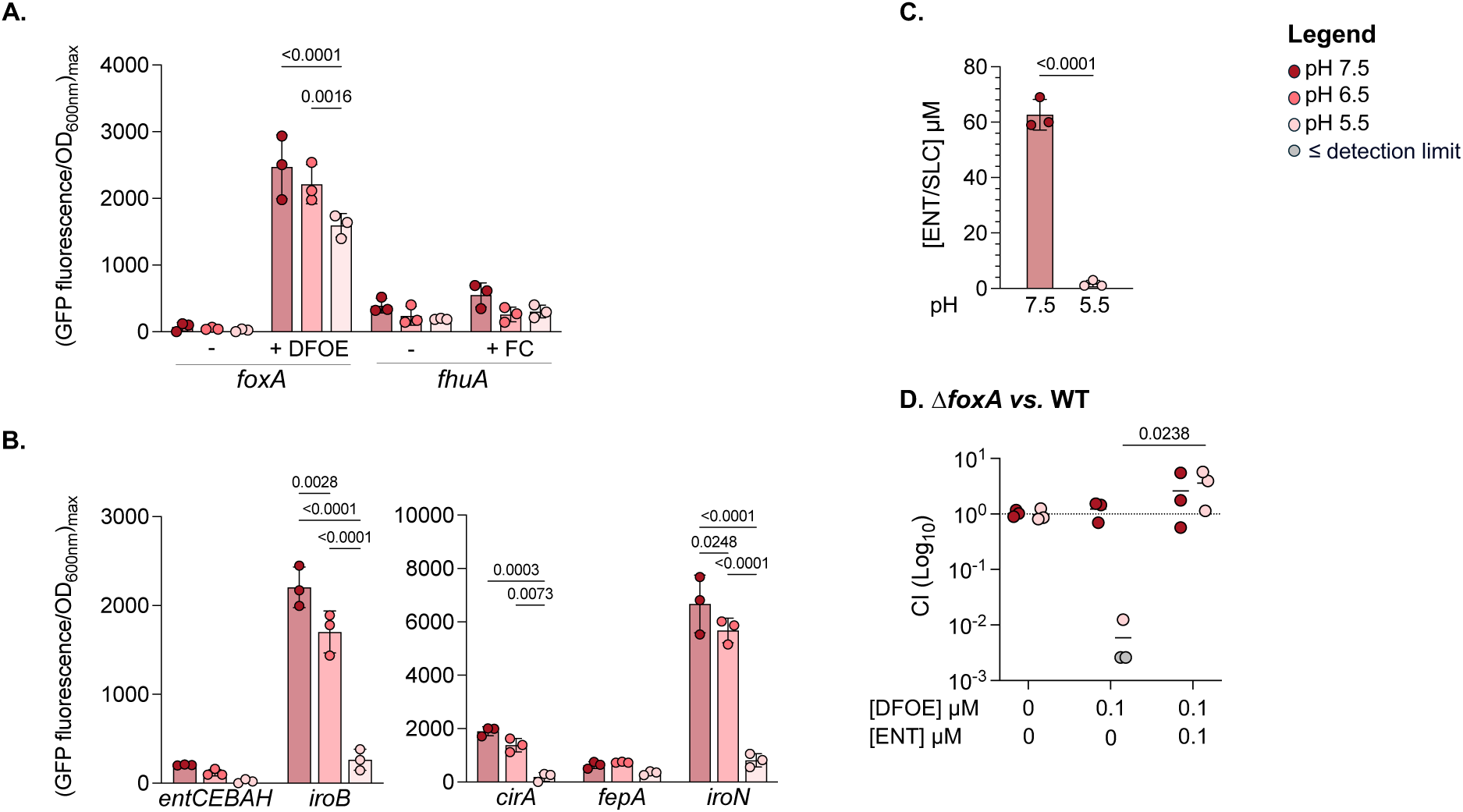
Endogenous siderophore production decreases in acidic conditions, heightening reliance on exogenous siderophores. **A-B**. Transcriptional expression of siderophore-related genes. WT strains carrying plasmid encoded fluorescent reporters were cultivated in MM at pH 7.5 (dark red), 6.5 (medium red), or 5.5 (light red). GFP-fluorescence emission data, normalized to optical density at 600 nm (OD_600nm_) of the culture, are shown as the maximum fluorescence per OD_600nm_. Each point represents a biological replicate. **A.** Expression levels of genes encoding exogenous siderophore transporters with or without 10 µM of their respective ligand. **B.** Expression levels of genes encoding transporters and enzymes involved in the biosynthesis of endogenous siderophores. **C**. Concentration of endogenous siderophores (ENT/SLC, and DHBS) in the supernatant of WT *Salmonella* cultures after 24 hours of cultivation in MM at pH 7.5 (dark red) and 5.5 (light red). The concentrations of ENT and SLC are shown together, as they have similar spectral properties, with SLC being a glycosylated derivative of ENT. **D.** Competitive indexes of Δ*foxA vs.* WT strain at pH 7.5 5 (dark red) and 5.5 (light red). The competition was conducted in MM with no supplementation, 0.1 μM DFOE, or 0.1 μM DFOE plus 0.1 μM ENT. The output ratio was calculated after 48 hours of culture. Each data point represents one biological replicate (n = 3). Points below the detection limit for each assay are shown in gray. Statistical analysis was performed using one-way ANOVA followed by Tukey’s multiple comparisons test (A, B, D) or unpaired t-test (C). Exact p-values are reported for all comparisons with p < 0.05 considered statistically significant.

### Loss of the sigma factor RpoS enhances endogenous siderophore production and acquisition at acidic pH

To elucidate the regulatory mechanisms underlying the reduced production of endogenous siderophores in acidic conditions, we investigated the roles of two potential candidates: Fur and RpoS (σ^70^). Fur is a key regulator of iron homeostasis in *Salmonella*, typically repressing iron import systems when iron is plentiful^46^. RpoS, a sigma factor, is involved in the transcription of genes associated with stress responses^64,65^. Given that our experimental conditions are both iron-poor and acidic, both Fur and RpoS were considered promising candidates for this study. Notably, the stress sigma factor RpoS has been shown to counteract Fur’s repression of genes involved in iron and manganese metabolism and modulate the metallome of *Salmonella enterica* serovar Typhimurium^66^. This suggests a potential regulatory interplay between Fur and RpoS in response to environmental stressors such as acidic conditions, where both factors could play a role in modulating iron homeostasis and siderophore production.

We measured endogenous siderophore concentrations in the supernatants of strains grown at pH 7.5 and 5.5 for both *fur* and *rpoS* mutant (Fig. 5A and fig. S5). The deletion of *fur* did not increase siderophore production. The fact that the depletion of Fur does not significantly impact siderophore production under our iron-limited conditions, suggests that in all our growth conditions Fur is completely de-repressed. While iron requirements may be reduced at low pH, *Salmonella* is still facing strict iron deprivation, highlighting the persistent iron limitation even under acidic stress. In contrast, the deletion of *rpoS* led to a marked increase in the concentration of endogenous siderophores in the growth medium, particularly at low pH, where the increase in ENT levels was significantly more pronounced than in the WT strain. At neutral pH, the difference was also evident, but the effect was far greater under acidic conditions (1.5-times more ENT at pH 7.5 versus 60-times more ENT at pH 5.5). Additionally, the *rpoS* mutant showed an enhanced iron deficiency response, as indicated by the increased expression of the small RNA *ryhB2* in all three pH conditions tested (Fig. 5B and fig. S5). This upregulation was also observed for genes related to both endogenous and exogenous siderophore import, such as *entCEBAH*, *fepA*, and *foxA* (Fig. 5C - 5E and fig. S5).

RpoS is known to modulate nutrient acquisition, typically by repressing the expression of various nutrient uptake systems. While at neutral pH iron-uptake systems are already active, the absence of RpoS has only limited effects on siderophore production. Contrary, at low pH, the absence of RpoS results in a more significant effect, where ENT production is strongly increased. This enhanced production of ENT at low pH underscores the stronger iron acquisition capacity in the *rpoS* mutant under these conditions. RpoS seems to have a significant role in regulating siderophore production under acidic conditions, whether directly or through intermediary pathways, leading to a drastic effect on growth under iron starvation.

**Figure 5.**
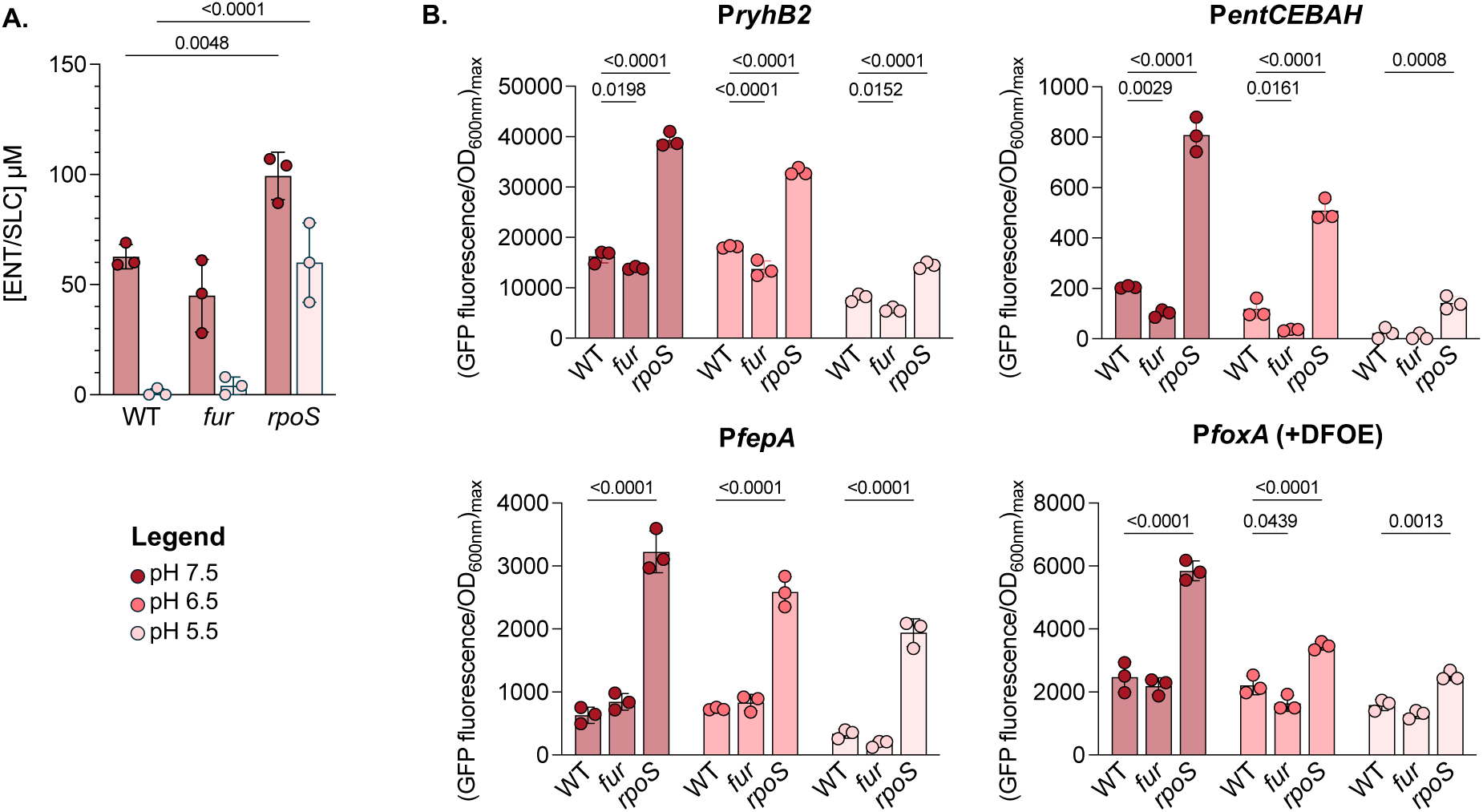
Loss of the sigma factor RpoS enhances endogenous siderophore production and acquisition at acidic pH. **A.** Measurement of endogenous siderophore production (ENT/SLC) in the Δ*fur* and Δ*rpoS* mutants grown in MM at pH 7.5 (dark red) and 5.5 (light red). Siderophore concentrations were quantified from the supernatant after 24 hours of cultivation (n = 3). **B-E.** Transcriptional expression of siderophore-related genes in Δ*fur* and Δ*rpoS* mutants. Strains were cultivated in MM at pH 7.5 (dark red), 6.5 (medium red), or 5.5 (light red). GFP-fluorescence emission data, normalized to optical density at 600 nm (OD_600nm_) of the culture, are shown as the maximum fluorescence per OD_600nm_. Each point represents one biological replicate (n = 3). **B**. Expression of the iron deprivation marker *ryhB2* (P*_ryhB2_*). **C.** Expression of the operon involved in endogenous siderophore biosynthesis *entCEBAH* (P*_entCEBAH_*). **D.** Expression of the transporter of ENT *fepA* (P*_fepA_*). **E.** Expression of the transporter of ferrioxamine E *foxA* (P*_foxA_*). Statistical analysis was performed using unpaired t-test (A) or one-way ANOVA followed by Tukey’s multiple comparisons test (B). Exact p-values are reported for all comparisons with p < 0.05 considered statistically significant.

## Discussion

For successful growth, organisms must adapt to their environments. Given the crucial role of iron for growth, the regulation of iron acquisition systems in bacteria must be finely tuned. This is especially critical for pathogens that need to establish infection in environments that are already colonized. This study explored how environmental changes, particularly a decrease in pH, influence the utilization of siderophores by *Salmonella*.

Our competition assays revealed that under acidic conditions, *Salmonella* benefits more from exploiting exogenous hydroxamate-type siderophores than it does at neutral pH, suggesting this as an adaptive strategy to acidic environments. This increased reliance on exogenous siderophores is linked to a stark reduction in the production of endogenous siderophores, monitored in various *Salmonella* strains (fig. S7), highlighting a drastic adaptation of siderophore production to varying pH changes. This phenomenon has been documented in previous studies. For instance, Valdebenito et *al*., demonstrated that in *Escherichia* coli, the production of ENT decreases under acidic conditions^48^. However, this strain, in contrast to our *Salmonella* strain, is also capable of producing other siderophores, such as aerobactin and yersiniabactin. Aerobactin, a mixed hydroxy acid and hydroxamate-type siderophore, reaches peak production at acidic pH, while yersiniabactin, a phenolate-type siderophore, is maximally produced at alkaline pH 8.5. This indicates that siderophore production in *E. coli* is highly influenced by environmental pH and varies with the type of siderophore^48^.

In addition to affecting siderophore production, environmental pH can also drive bacterial adaptation at the level of siderophore transport. Li et *al*., found that *Acinetobacter baumannii* adapts to pH changes through mutations in the siderophore receptor gene *bauA*^51^. This adaptation allows *A. baumannii* to optimize iron uptake and maintain iron levels under varying pH conditions. To explore whether a similar pH-dependent adaptation occurs in *Salmonella*, we investigated how pH affects the affinity of ferrioxamine for its transporter FoxA. However, our results revealed no significant change in ferrioxamine binding to FoxA across different pH levels (Fig. S4).

Not only can pH affect the affinity of siderophores for their transporter, but it also influences their affinity for iron by modifying their chelating properties. ENT, a catecholate-type siderophore, is known to have the highest iron affinity at neutral pH (pFe at pH 7.5 = 35) among all the siderophores described^67^. However, Valdebenito et *al*., found that this affinity decreases under acidic conditions (pFe at pH 5.5 = 25). In contrast, hydroxamate siderophores like deferoxamine maintain relatively stable iron affinities across different pH levels (pFe-pH 7.5 = 27 and pFe pH-5.5 = 22)^48^. Our competition assays, where ENT and DFOE were added simultaneously at equal concentration, revealed that *Salmonella* favors ENT over DFOE even under acidic conditions. The superior iron-binding capacity of ENT makes it the preferred siderophore when both are available, underlining yet again that the stark dependency on exogenous siderophore at acidic pH is solely due to reduced ENT/SLC production.

Upon exposure to low pH, *Salmonella* activates the alternative sigma factor RpoS, which orchestrates the acid tolerance response by upregulating the expression of numerous acid shock proteins but also altering nutrient acquisition, thereby enabling the pathogen to survive in acidic environments^61^ . Our data suggest that in acidic environments, RpoS but not Fur reduces siderophore production and thereby increases *Salmonella’s* reliance to exogenous siderophores. The exploitation of exogenous siderophores may allow *Salmonella* to avoid the energy costs associated with producing its own siderophores. This emphasis on energy conservation likely explains why the deletion of *rpoS* positively impacts endogenous siderophore production. The loss of RpoS has already been shown to enhance *Salmonella*’s ability to scavenge for limited nutrients^68^. While this strategy may be less critical at neutral pH, it may become especially important under acidic conditions, where conserving energy is vital for enduring the environmental stress and represent an evolutive response to stress conditions.

Understanding the conditions under which *Salmonella* utilizes endogenous or exogenous siderophores can provide critical insights for designing targeted therapies. Despite the challenges in pinpointing specific targets, knowing how *Salmonella* adapts its iron acquisition strategies offers valuable information for developing effective treatments. Future research should focus on exploring other environmental clues that could affect iron acquisition mechanisms and thereby offering a better understanding versatility of *Salmonella’s* iron uptake strategies and thereby informing future therapeutic approaches.

## Acknowledgments

We are indebted to members of the MMBCA lab for discussion. We thank Prof. Kevin Foster for sharing *Salmonella* strains.

## Author contributions

Conceptualisation: OC. Methodology: MF, CS and OC. Investigation: MF and OC. Visualisation: MF and OC. Funding acquisition: OC. Supervision: OC, CS and IJS. Writing – original draft: MF and OC. Writing – review and editing: MF, CS, IJS and OC. ChatGPT version 4.o was sometimes used for Language improvement of the manuscript.

## Data and materials availability

All data are available in the main text or supplementary materials and on https://doi.org/10.5281/zenodo.15276328.

## Materials and methods

### Bacterial strains and growth conditions

A full list of all bacterial strains used in this study is provided in sup. table S1. *Salmonella* strains used were based on sME51, a prototrophic *hisG*^Leu69^ derivative of *Salmonella enterica* serovar Typhimurium SL1344^69^. All strains were cultivated in Lysogeny Broth (LB) (10 g/L tryptone, 5 g/L yeast extract, 5 g/L NaCl) or Minimal Medium (MM) (15 mM NH_4_Cl, 1.5 mM K_2_SO_4_, 3 mM KH_2_PO_4_, 1 mM MgCl_2_, 0.4% glycerol). For pH adjustments, 100 mM HEPES was used to achieve pH 7.5, while 100 mM MES was used for pH 6.5 and 5.5 and filter-sterilized. Media were supplemented with 90 µg/ml streptomycin, and 50 µg/ml kanamycin was added when a plasmid reporter was present. *Escherichia coli* JKe201 was grown in LB supplemented with 100 µM diaminopimelic acid (DAP), and 50 µg/ml kanamycin was added when required. Cultures were incubated at 37°C with shaking at 220 rpm.

### Plasmids and genetic engineering of bacterial strains

All primers and plasmids used in this study are listed in sup. table S2 and S3. All plasmids used in this study are low-copy plasmids and carried a kanamycin resistance cassette to ensure their maintenance in bacterial cells. Gene reporter plasmids contained candidate gene promoters fused to a *gfp-ova* reporter to assess specific gene expression in response to experimental conditions as described here^59,60^. For strain differentiation during competition assays, low-copy episomal pSC101-derivatives were employed. These plasmids enabled the constitutive expression of either dTomato or YPet from the P*_ybaJ_* promoter, which facilitated tracking the proportions of different strains^59,69^. Gene deletions were obtain using two consecutive single-crossovers, incorporating positive selection for kanamycin resistance and a double negative selection against SacB levan sucrase-mediated sensitivity to sucrose and the expression of the I-SceI endonuclease^70^. The engineered strains preserved a nonapeptide with the first five and last four amino acids, including the stop codon, as described here^70^.

### Prevalence of siderophore-related genes and phylogenetic analysis in *Salmonella*

To analyze siderophore transporter genes across *Salmonella* strains, we followed a structured bioinformatics workflow. Initially, we sorted and downloaded genomes from the BV-BRC database^55^. Genomes were first assessed for contamination and metadata. We then filtered these genomes to retain only those with a minimum size of 4 million base pairs. To ensure the most representative dataset, we grouped strains by their unique multilocus sequence typing (MLST) type and randomly selected one strain per group using Python’s *random.choice ()* function. This approach allowed us to capture the diversity of *Salmonella* strains effectively and resulted in a final dataset of 183 strains, selected from an initial set of 1,146 isolates. For gene detection, we used the ABRicate tool^71^ with a custom database to identify specific siderophore-related genes in the genome sequences. Genes were considered present only if they met the default ABRicate thresolds of at least 75% identity and 60% coverage; any gene below these thresholds was classified as absent. Phylogenetic analysis was conducted using PhyloPhlAn^72^. We first generated a database of core proteins for *Salmonella* and then reconstructed the phylogenetic tree by trimming sequences, filtering data, and using FastTree and RAxML for tree construction. Maximum likelihood trees were inferred using concatenated gene sequences with the GTR model and bootstrap support. The resulting tree was visualized and further modified using iTOL (version 7.0) to enhance clarity and presentation.

### Competition assays

The competing strains carried a low-plasmid with constitutive expression of fluorescent proteins (YPet or dTomato). Pre-cultures were grown overnight in MM at pH 7.5 with 10 μM iron, at 37°C with agitation (220 rpm). Post-incubation, 1 mL of each culture was centrifuged at 9000 rpm for 2 minutes, washed twice with 1X phosphate-buffered saline (PBS), and resuspended to an optical density at 600 nm of 0.1 in 1 mL of 1X PBS. Equal volumes (500 µL each) of the two strains were mixed to create a 1:1 bacterial suspension at OD_600nm_ of 0.1 (OD 0.05 per strain). Fourteen-milliliter tubes containing 1.8 mL of MM at pH 7.5, 6.5, or 5.5, with or without exogenous siderophore (all chemicals are reported in sup. table S4), were inoculated with 200 µL of the bacterial mixture to achieve a final OD_600nm_ of 10^-6^. Cultures were grown for 48 hours at 37°C with agitation (220 rpm). The total number of cells for each strain was determined by plating on LB agar supplemented with 90 μg/mL streptomycin and enumerating colonies under blue light to distinguish the fluorescent proteins. The competitiveness index (CI) was calculated as CI = output/input, where output is the ratio of mutant to wild-type colonies after 48 hours, and input is the ratio at time zero.

### Quantification of GFP-fluorescence intensity

Overnight cultures in MM at pH 7.5 with 10 μM iron were grown at 37°C with shaking (220 rpm). Following centrifugation (9000 rpm, 2 minutes) and washing twice with 1X PBS, cells were resuspended to an OD_600nm_ of 0.1 in 1X PBS. Twenty microliters of this suspension were added to 180 μL of fresh MM at various pH levels (7.5, 6.5, 5.5) with or without iron or exogenous siderophore in a 96-well plate (Greiner, U-bottomed microplate). The plate was incubated at 37°C with shaking in a Tecan microplate reader (Infinite M200, Tecan), with OD_600nm_, GFP (excitation/emission: 488 nm/510 nm), and mCherry (excitation/emission: 570 nm/610 nm) fluorescence measured every 10 minutes for 48 hours. Results from three biological replicates were analyzed, with GFP fluorescence normalized to OD_600nm_ and the maximum value of this normalization used for histograms. Statistical significance was determined with p-values <0.05.

### Ligand binding assay

To determine the dissociation constant (Kd) between Ferri-DFOE and its transporter FoxA in *Salmonella*, strains were cultured overnight in MM at pH 7.5 or 5.5, with 10 μM DFOE added to induce FoxA transporter expression. Complexes of ^55^FeCl_3_-DFOE were prepared with a ^55^Fe concentration of 20 μM and a siderophore ratio of 20:1. A known concentration of *Salmonella* was incubated with ^55^FeCl_3_-DFOE complexes in cold-buffer at pH 7.5 (100 mM HEPES) or pH 5.5 (100 mM MES) for 1 hour at 0°C, with transport inhibited by adding Carbonyl cyanide 3-chlorophenylhydrazone. Bacterial cells bound to the complexes were recovered by filtration using a Brandel filtration system (M-48 Cell Harvester) and Whatman® glass microfiber filters (Grade GF/B, 460x570 mm). Radioactivity associated with the bacteria was measured with a scintillation counter to calculate the K*d*. This binding assay was conducted in triplicate.

### Siderophore production

Overnight cultures grown in MM at pH 7.5 with 10 μM iron were pelleted, re-suspended in fresh MM at pH 7.5 or 5.5 to an OD_600nm_ of 0.01, and incubated with shaking at 37°C for 24 hours. The OD_600nm_ was measured to assess cell growth. Supernatants were analyzed by UV-Vis spectroscopy. Preliminary, calibration curves with pure ENT and its degradation product, DHBS, were established, with absorbance peaks at 330 nm for ENT and 310 nm for DHBS at pH 7.5, and at 317 nm for ENT and 310 nm for DHBS at pH 5.5. Molar extinction coefficients were εENT pH 7.5 = 8100 M^-1^.cm^-1^, εENT pH 5.5 = 2290 M^-1^.cm^-1^, εDHBS pH 7.5 = 7000 M^-1^.cm^-1^, and εDHBS pH 5.5 = 2100 M^-1^.cm^-1^. Absorbance monitored from supernatants were normalized to OD_600nm_ of the cultures. Control experiments compared the absorbance of ENT/DHBS in wild-type strains with those deleted for siderophore production to account for interference from other media components.

### Statistics

Statistical tests were performed with GraphPad Prism (version 10.2.3) as indicated in the figure legends. Figure legends indicate the statistical tests used with all biological replicates depicted in the figures.

## Supplementary figures

**Fig. S1.**
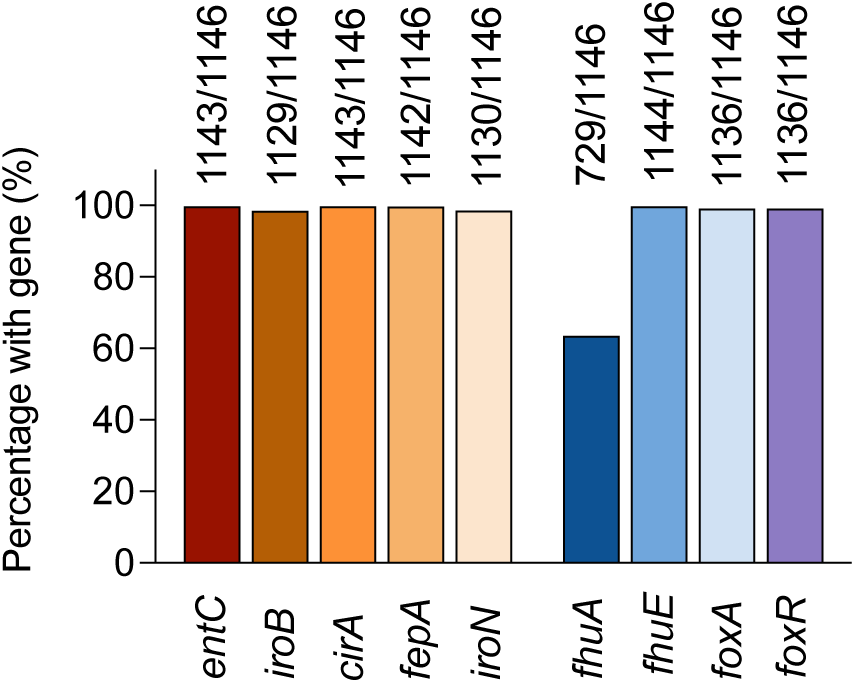
Conservation of siderophore-related genes among 1,146 *Salmonella* strains from the BV-BRC database. Bars in warm colours indicate the proportion of tested strains carrying genes involved in endogenous siderophore biosynthesis (*entC*, *iroB*) and uptake (*cirA*, *fepA*, *iroN*). Bars in cool colors indicate the proportion of tested strains carrying genes involved in exogenous siderophore uptake (*fhuA*, *fhuE* and *foxA*) and its regulation (*foxR*). These isolates were selected following a structured bioinformatics workflow, which included genome quality assessment, contamination checks, and filtering based on a minimum genome size of 4 million base pairs (see Methods).

**Fig. S2.**
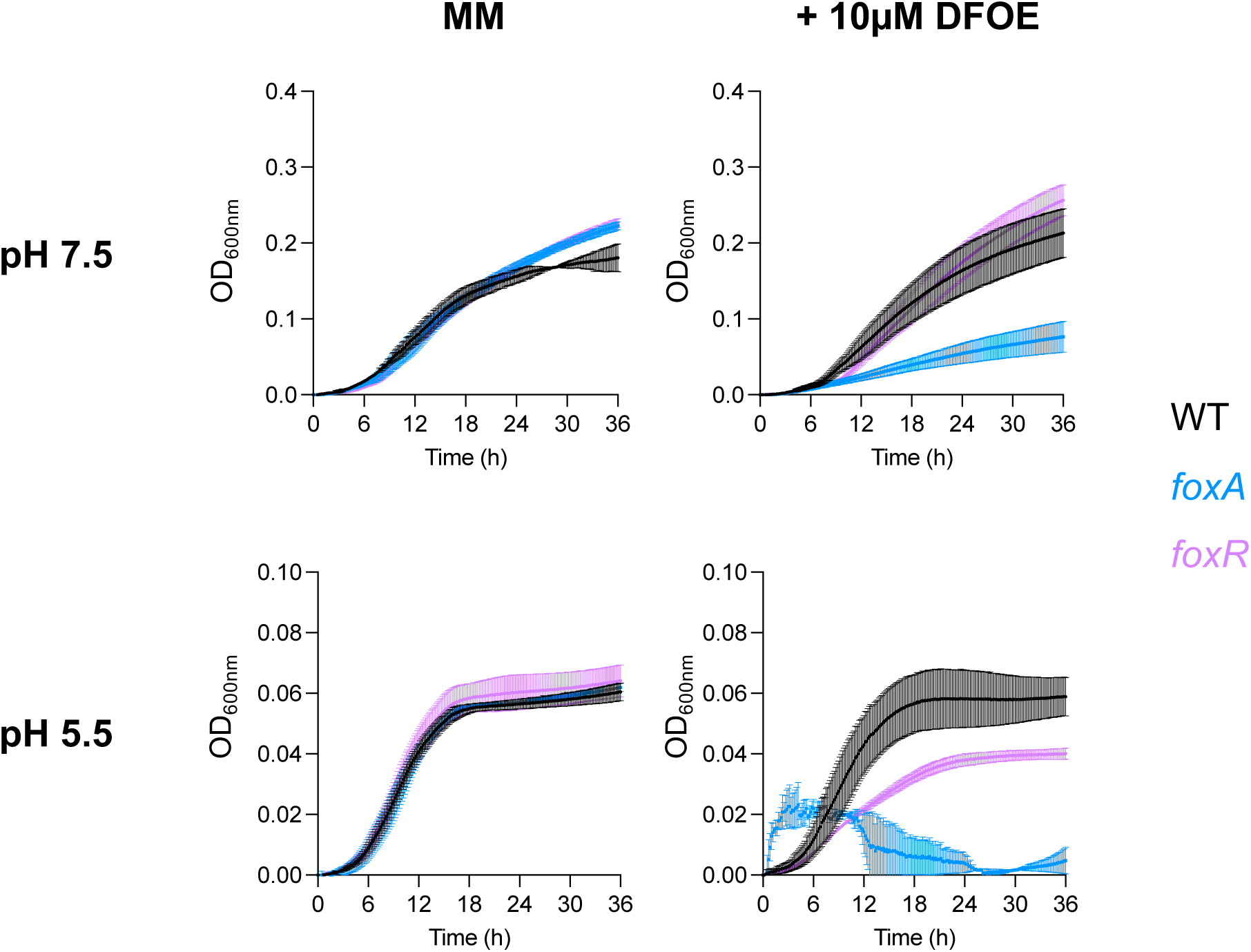
*Salmonella* WT DFOE-dependent growth kinetics at various pH (7.5 vs 5.5). Bacterial strains were cultivated in iron-depleted medium at either neutral (pH 7.5) or acidic (pH 5.5) conditions, in the absence or presence of 10 µM of DFOE. Growth was initiated at an OD_600nm_ of 0.01 and monitored over time. The WT strain (black) was compared to the *foxA* deletion mutant (blue) which cannot exploit DFOE, and the *foxR* deletion mutant (purple), which is unable to regulate *foxA* expression in response to DFOE. Each growth curve represents the median with the standard error of mean of three biological replicates.

**Fig. S3.**
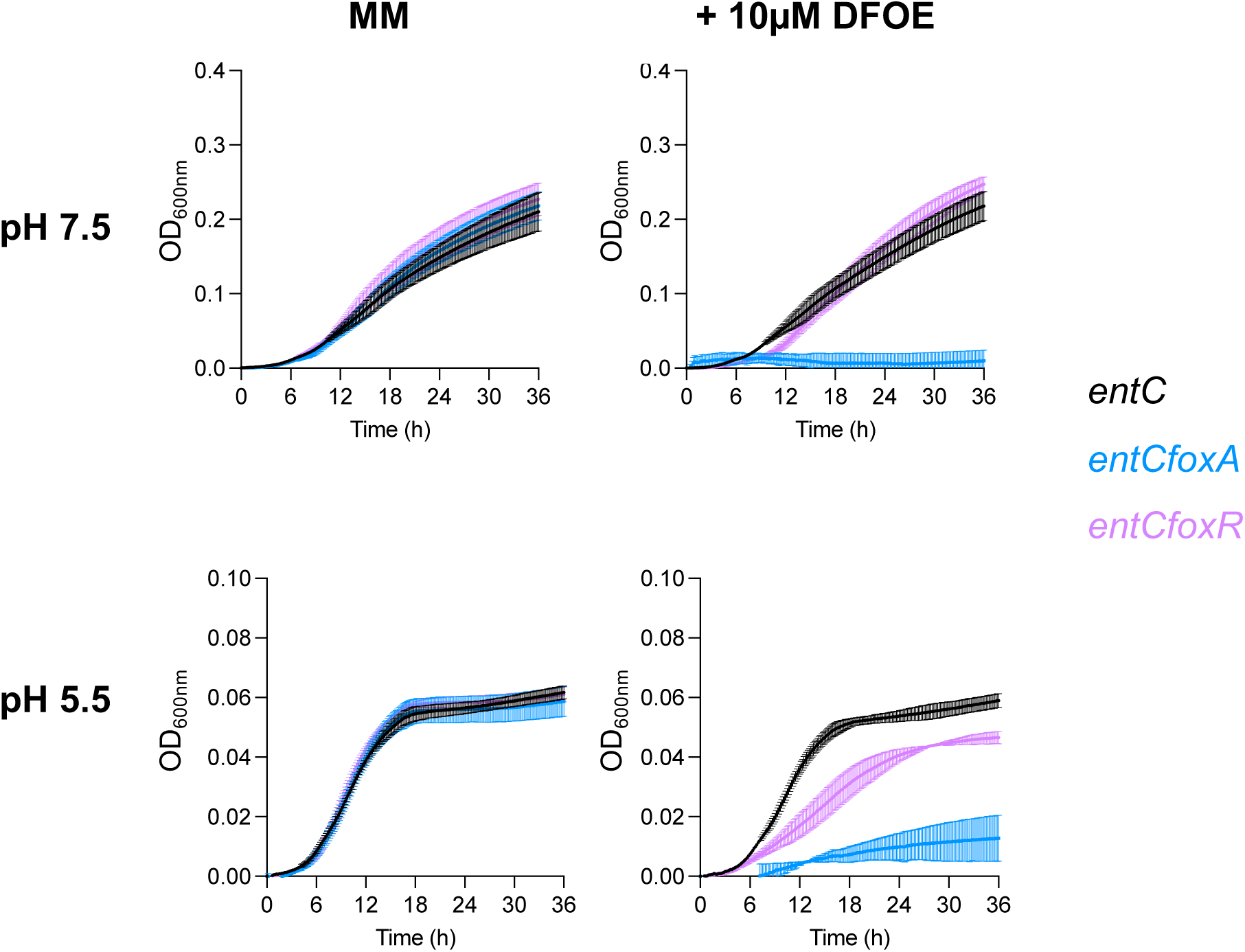
*Salmonella entC* mutant DFOE-dependent growth kinetics at various pH (7.5 vs 5.5). Bacterial strains were cultivated in iron-depleted medium at either neutral (pH 7.5) or acidic (pH 5.5) conditions, in the absence or presence of 10 µM of DFOE. Growth was initiated at an OD_600nm_ of 0.01 and monitored over time. The *entC* mutant (black), which cannot produce endogenous siderophores but can exploit DFOE, was compared to the *entCfoxA* double mutant (blue), which lacks both endogenous siderophore production and the ability to exploit DFOE, and the *entCfoxR* double mutant (purple), which lacks endogenous siderophore production and cannot regulate *foxA* expression in response to DFOE. Each growth curve represents the median with the standard error of mean of three biological replicates.

**Fig. S4.**
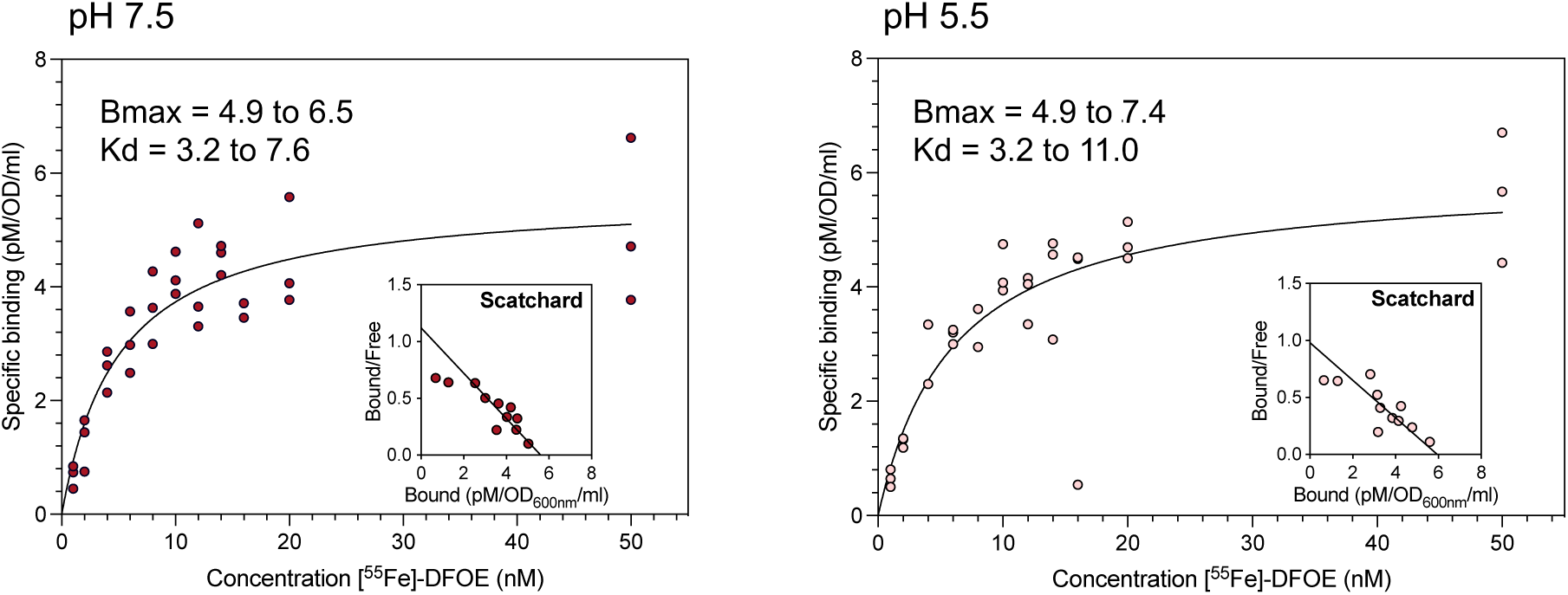
Binding of Fe-DFOE to FoxA in *Salmonella* in function of the pH (7.5 vs 5.5). Bacterial cells were incubated with a series of concentrations of [^55^Fe]-DFOE at 0°C for 1 hour in HEPES buffer (pH 7.5) or MES buffer (pH 5.5), with transport inhibited by carbonyl cyanide 3-chlorophenylhydrazone (CCCP). [^55^Fe]-DFOE binding was assessed by filtration and radioactivity associated with the cells was measured by scintillation counting to determine the dissociation constant (K*d*) and the maximum binding capacity (Bmax). The saturation binding curves represent the specific binding of Fe-DFOE to FoxA at both pH conditions. Inset, Scatchard analysis of specific binding. Each data point represents one biological replicate (n = 3).

**Fig. S5.**
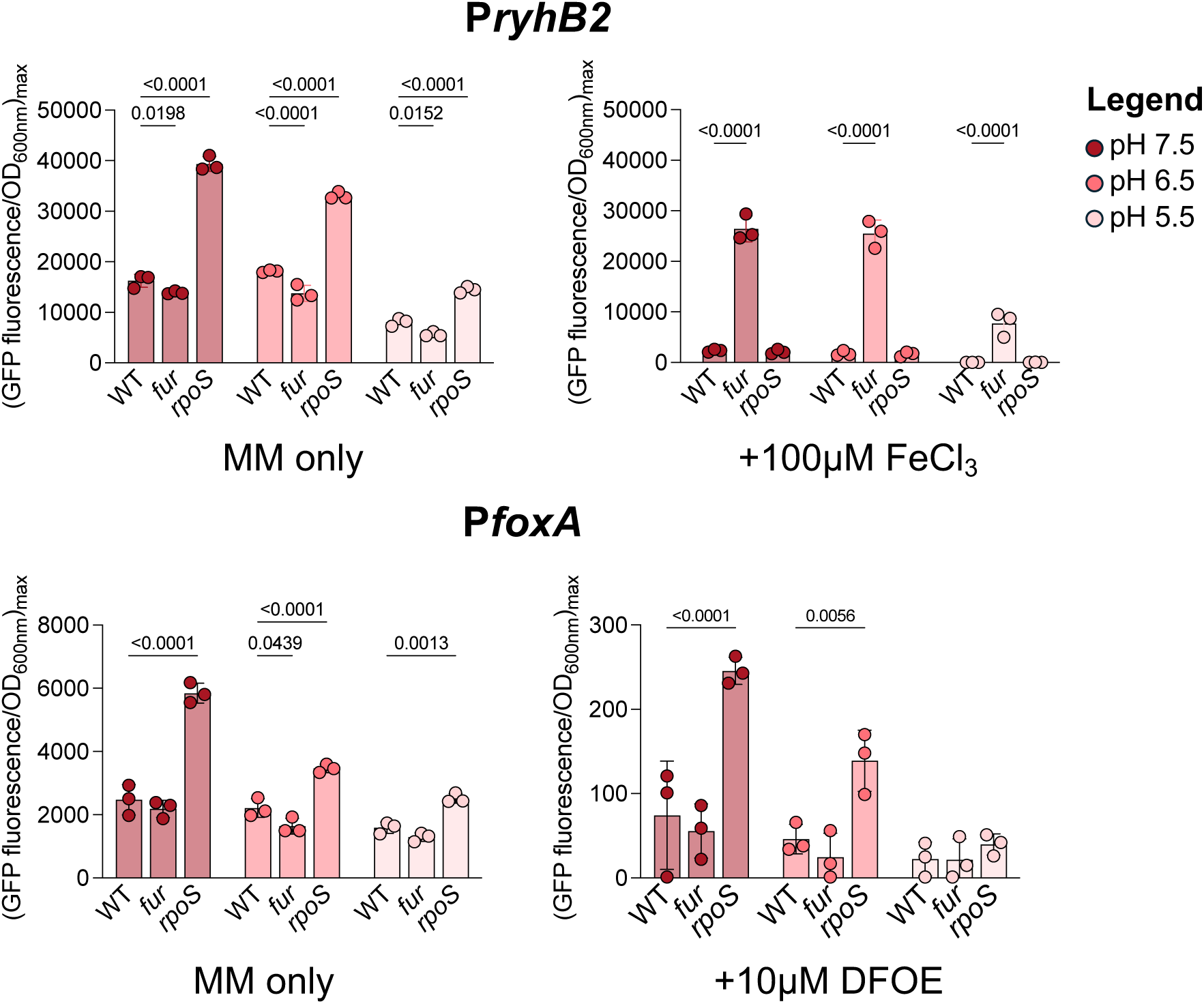
Transcriptional expression of *ryhB2* (P*_ryhB2_*) and *foxA* (P*_foxA_*) in *Salmonella* at different pH levels, in iron-poor and iron-supplemented medium. Strains were cultivated in MM under iron-depleted conditions or supplemented with 100 µM FeCl_3_ (PryhB2) or 10 µM DFOE (PfoxA). Cultures were grown at pH 7.5 (dark red), pH 6.5 (medium red), and pH 5.5 (light red). Fluorescence emission from P*ryhB2*-*gfp* and P*foxA*-*gfp* transcriptional reporters, normalized to OD_600nm_. The maximum fluorescence signal during growth is shown. Expression was compared between the WT strain, a *fur* deletion mutant and a *rpoS* deletion mutant. Each point represents one biological replicate (n = 3). Statistical analysis was performed using one-way ANOVA followed by Tukey’s multiple comparisons test. Exact p-values are reported for all comparisons with p < 0.05 considered statistically significant.

**Fig. S6.**
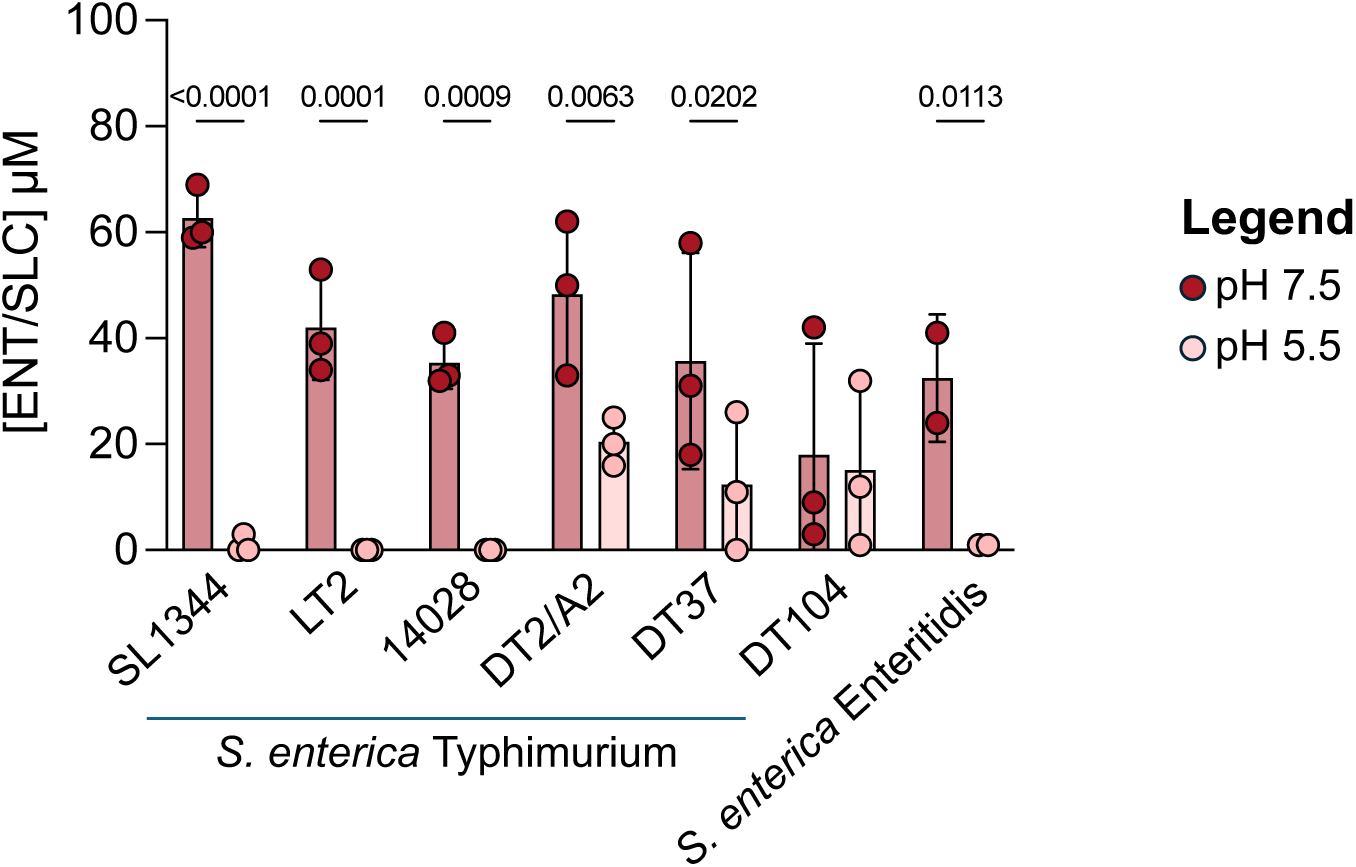
Endogenous siderophore production (ENT/SLC) by different strains of *S. enterica* in function of pH. Endogenous siderophore concentrations were measured in the supernatant of various *S. enterica* strains after 24 hours of cultivation in MM under iron-depleted at pH 7.5 (dark red) and pH 5.5 (light red). The concentrations of ENT and SLC are shown together as SLC is a glycosylated derivative of ENT. Each point represents one biological replicate (n = 3). Statistical analysis was performed using an unpaired t-test.

**Fig. S7.**
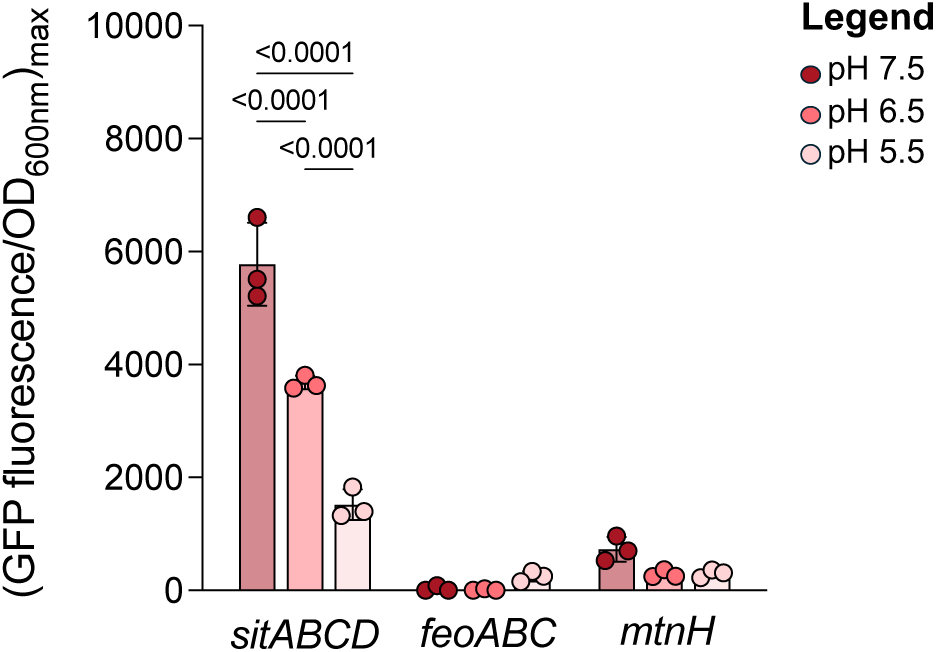
Transcriptional expression of the ferrous iron uptake systems in *Salmonella* (P*_sitABCD_*, P*_feoABC_* and P*_mntH_*) at different pH levels. Strains were cultivated in MM under iron-depleted at pH 7.5 (dark red), pH 6.5 (medium red), and pH 5.5 (light red). Fluorescence emission from P*sitABCD*-*gfp*, P*feoABC*-*gfp* and P*mntH*-*gfp* transcriptional reporters was normalized to OD_600nm_. The maximum fluorescence signal during growth is shown. Each point represents one biological replicate (n = 3). Statistical analysis was performed using one-way ANOVA followed by Tukey’s multiple comparisons test. Exact p-values are reported for all comparisons with p < 0.05 considered statistically significant.

## Figure captions

Tab. S1. Strains used in this study.

Tab. S2. Primers used in this study.

Tab. S3. Plasmids used in this study.

Tab. S4. Reagents used in this study.

Tab. S5. Strain information and gene blast results of the 1,186 strains of *Salmonella*. Found on https://doi.org/10.5281/zenodo.15276328

