## Supplemental tables for "*Salmonella* relies on siderophore exploitation at low pH"

### Supplementary table S1: strains used in this study.

| ***Escherichia coli*** | | |
|---|---|---|
| **Strain** | **Description** | **Source** |
| JKe201 | MG1655 RP4-2-Tc::[ΔMu1::Δ*aac(3)IV*::*lacIq*-Δ*aphA*-Δ*nic35*-ΔMu2::*zeo*] Δ*dapA*::(*erm-pir*) Δ*recA* Δ*mcrA* Δ(*mrr-hsdRMS-mcrBC*) | (1) |
| ***Salmonella enterica*** | | |
| **Strain** | **Phenotype** | **Source** |
| SL1344 *hisGLeu69* | HisG prototrophe *Salmonella enterica* SL1344 strain used as WT | (2), (3) |
| *entC* | SL1344 *hisGLeu69* - deletion in *entC* gene | (4) |
| *foxA* | SL1344 *hisGLeu69* - deletion in *foxA* gene | This study |
| *fhuA* | SL1344 *hisGLeu69* - deletion in *fhuA* gene | This study |
| *fhuE* | SL1344 *hisGLeu69* - deletion in *fhuE* gene | This study |
| *foxR* | SL1344 *hisGLeu69* - deletion in *foxR* gene | This study |
| *entC foxA* | SL1344 *hisGLeu69* - deletion in *entC* and *foxA* genes | This study |
| *entC foxR* | SL1344 *hisGLeu69* - deletion in *entC* and *fhuA* genes | This study |
| *feoABC sitABCD mntH* | SL1344 *hisGLeu69* - deletion in *feoABC, sitABCD* and *mntH* genes | (4) |
| *fur* | SL1344 *hisGLeu69* - deletion in *fur* gene | This study |
| *rpoS* | SL1344 *hisGLeu69* - deletion in *rpoS* gene | This study |
| *fur rpoS* | SL1344 *hisGLeu69* - deletion in *fur* and *rpoS* genes | This study |
| LT2 | *Salmonella enterica enterica* serovar Typhimurium LT2 | *K. Foster lab* |
| 14028S | *Salmonella enterica enterica* serovar Typhimurium 14028S | *K. Foster lab* |
| DT2/A2 | *Salmonella enterica enterica* serovar Typhimurium DT2/A2 | *K. Foster lab* |
| DT37 | *Salmonella enterica enterica* serovar Typhimurium DT37 | *K. Foster lab* |
| DT104 | *Salmonella enterica enterica* serovar Typhimurium DT104 | *K. Foster lab* |


[1]A. Harms, M. Liesch, J. Körner, M. Québatte, P. Engel, and C. Dehio, “A bacterial toxin-antitoxin module is the origin of inter-bacterial and inter-kingdom effectors of Bartonella,” PLoS Genet, vol. 13, no. 10, p. e1007077, Oct. 2017, doi: 10.1371/journal.pgen.1007077.

[2]S. K. Hoiseth and B. A. Stocker, “Aromatic-dependent Salmonella typhimurium are non-virulent and effective as live vaccines,” Nature, vol. 291, no. 5812, pp. 238–239, May 1981, doi: 10.1038/291238a0.

[3]B. Claudi et al., “Phenotypic variation of Salmonella in host tissues delays eradication by antimicrobial chemotherapy,” Cell, vol. 158, no. 4, pp. 722–733, Aug. 2014, doi: 10.1016/j.cell.2014.06.045.

[4]O. Cunrath and D. Bumann, “Host resistance factor SLC11A1 restricts Salmonella growth through magnesium deprivation,” Science, vol. 366, no. 6468, pp. 995–999, Nov. 2019, doi: 10.1126/science.aax7898.

[5]F. R. Cianfanelli, O. Cunrath, and D. Bumann, “Efficient dual-negative selection for bacterial genome editing,” BMC Microbiol, vol. 20, no. 1, p. 129, May 2020, doi: 10.1186/s12866-020-01819-2.

[6]B. Roche et al., “Heterogeneous dual-metal control of Salmonella infection,” Oct. 24, 2023, bioRxiv. doi: 10.1101/2023.10.23.562652.

#### Supplementary table S2: primers used in this study.

| Primer name | Sequence | Description |
|---|---|---|
| oOPC-118 | CCCAGTCTCGAGGTCGACGGTATCGATAAGCTTGATATCGAATTCagccaggtgactatcacctgggtaggc | amplify 700bp downstream of *fhuA* for pOPC-32 for *fhuA* deletion in *Salmonella enterica* SL1344 |
| oOPC-119 | agaaattagaaacggaaggttttaagacgcgccattggtatatct | |
| oOPC-120 | taccaatggcgcgtcttaaaaccttccgtttctaatttctttttgggcacg | amplify 700bp upstream of *fhuA* for pOPC-32 for *fhuA* deletion in *Salmonella enterica* SL1344 |
| oOPC-121 | CTGGAGCTCCACCGCGGTGGCGGCCGCTCTAGAACTAGTGGATCCgggccgccatgttgatatcgtgcag | |
| oOPC-402 | acaatgtcgatacctggtttgccgg | check *fhuA* deletion in *Salmonella enterica* SL1344 |
| oOPC-403 | cgccgcgtagcgcgactaaataatc | |
| oOPC-955 | CCCAGTCTCGAGGTCGACGGTATCGATAAGCTTGATATCGAATTCcgagcgctggttcacgctgcc | amplify 700bp downstream of *foxR* for pOPC-227 for *foxR* deletion in *Salmonella enterica* SL1344 |
| oOPC-956 | gcgcgttacgcacagtggcgggagatcggagacattatcaatacc | |
| oOPC-957 | tgataatgtctccgatctcccgccactgtgcg | amplify 700bp upstream of *foxR* for pOPC-227 for *foxR* deletion in *Salmonella enterica* SL1344 |
| oOPC-958 | CTGGAGCTCCACCGCGGTGGCGGCCGCTCTAGAACTAGTGGATCCcagcttcacttctcctccggcatca | |
| oOPC-959 | cgctcacccacgacgacatc | check *foxR* deletion in *Salmonella enterica* SL1344 |
| oOPC-960 | cgctggttgttattgccggc | |
| oOPC-156 | CCCAGTCTCGAGGTCGACGGTATCGATAAGCTTGATATCGAATTCtgagcggaatgcctgccag | amplify 700bp downstream of *fur* for pOPC-39 for *fur* deletion in *Salmonella enterica* SL1344 |
| oOPC-157 | tccgcatgactgacaacaatgacgcgactaaataagtgtaaatctttcgaagagc | |
| oOPC-158 | tacacttatttagtcgcgtcattgttgtcagtcatgcggaatctgtcc | amplify 700bp upstream of *fur* for pOPC-39 for *fur* deletion in *Salmonella enterica* SL1344 |
| oOPC-159 | CTGGAGCTCCACCGCGGTGGCGGCCGCTCTAGAACTAGTGGATCCacctggaagggtatgacattctg | |
| oMF-67 | CCCAGTCTCGAGGTCGACGGTATCGATAAGCTTGATATCGcggcttgctggccatca | amplify 700bp downstream of *rpoS* for pMF-12 for *rpoS* deletion in *Salmonella enterica* SL1344 |
| oMF-68 | accttatgagtcagaatacgctgttccgcgagtaagt | |
| oMF-69 | ggtacttactcgcggaacagcgtattctgactcataaggtgg | amplify 700bp upstream of *rpoS* for pMF-12 for *rpoS* deletion in *Salmonella enterica* SL1344 |
| oMF-70 | CTGGAGCTCCACCGCGGTGGCGGCCGCTCTAGAACTAGTGttactggcggaaatgcgatcacc | |
| oMF-71 | tttgggtatgccgtggacgg | check *rpoS* deletion in *Salmonella enterica* SL1344 |
| oMF-72 | aaccggtcacggaacaaccc | |

### Supplementary table S3: plasmids used in this study.

| Plasmid name | Relevant genotype | Resistance | Source |
|---|---|---|---|
| pFOK | *ori* R6K *y, KanR, I-SceI site, oriT, traJ, tetR, PtetA-I-SceI-sacB* | Kanamycin | (5) |
| pOPC-63 | pFOK-derived suicide vector to delete *entC* in *S. enterica* | Kanamycin | (4) |
| pOPC-175 | pFOK-derived suicide vector to delete *foxA* in *S. enterica* | Kanamycin | (5) |
| pOPC-32 | pFOK-derived suicide vector to delete *fhuA* in *S. enterica* | Kanamycin | This study |
| pOPC-227 | pFOK-derived suicide vector to delete *foxR* in *S. enterica* | Kanamycin | This study |
| pOPC-39 | pFOK-derived suicide vector to delete *fur* in *S. enterica* | Kanamycin | This study |
| pMF-12 | pFOK-derived suicide vector to delete *rpoS* in *S. enterica* | Kanamycin | This study |
| pOPC-176 | pFOK-derived suicide vector to delete *sitABCD* in *S. enterica* | Kanamycin | (5) |
| pOPC-225 | pFOK-derived suicide vector to delete *feoABC* in *S. enterica* | Kanamycin | (4) |
| pOPC-107 | pFOK-derived suicide vector to delete *mntH* in *S. enterica* | Kanamycin | (4) |
| pOPC-57 | R-$P^{foxA}$-*gfp* ova-$P^{ybaJ}$-*mCherry*-SC101 ori-*rep101*-*aphT* | Kanamycin | (6) |
| pOPC-55 | R-$P^{fhuA}$-*gfp* ova-$P^{ybaJ}$-*mCherry*-SC101 ori-*rep101*-*aphT* | Kanamycin | (6) |
| pOPC-20 | R-$P^{entcebah}$-*gfp* ova-$P^{ybaJ}$-*mCherry*-SC101 ori-*rep101*-*aphT* | Kanamycin | (6) |
| pOPC-213 | R-$P^{iroB}$-*gfp* ova-$P^{ybaJ}$-*mCherry*-SC101 ori-*rep101*-*aphT* | Kanamycin | (6) |
| pOPC-53 | R-$P^{cirA}$-*gfp* ova-$P^{ybaJ}$-*mCherry*-SC101 ori-*rep101*-*aphT* | Kanamycin | (6) |
| pOPC-54 | R-$P^{fepA}$-*gfp* ova-$P^{ybaJ}$-*mCherry*-SC101 ori-*rep101*-*aphT* | Kanamycin | (6) |
| pOPC-56 | R-$P^{iroN}$-*gfp* ova-$P^{ybaJ}$-*mCherry*-SC101 ori-*rep101*-*aphT* | Kanamycin | (6) |
| pOPC-18 | R-$P^{ryhB2}$-*gfp* ova-$P^{ybaJ}$-*mCherry*-SC101 ori-*rep101*-*aphT* | Kanamycin | (6) |
| pBC-20 | $P^{ybaJ}$-*Ypet*-SC101 ori-*rep101* -*aphT* | Kanamycin | (3) |
| pBC-53 | $P^{ybaJ}$-*dTomato*-SC101 ori-*rep101* -*aphT* | Kanamycin | (3) |

[1]A. Harms, M. Liesch, J. Körner, M. Québatte, P. Engel, and C. Dehio, “A bacterial toxin-antitoxin module is the origin of inter-bacterial and inter-kingdom effectors of Bartonella,” PLoS Genet, vol. 13, no. 10, p. e1007077, Oct. 2017, doi: 10.1371/journal.pgen.1007077.
[2]S. K. Hoiseth and B. A. Stocker, “Aromatic-dependent Salmonella typhimurium are non-virulent and effective as live vaccines,” Nature, vol. 291, no. 5812, pp. 238–239, May 1981, doi: 10.1038/291238a0.
[3]B. Claudi et al., “Phenotypic variation of Salmonella in host tissues delays eradication by antimicrobial chemotherapy,” Cell, vol. 158, no. 4, pp. 722–733, Aug. 2014, doi: 10.1016/j.cell.2014.06.045.
[4]O. Cunrath and D. Bumann, “Host resistance factor SLC11A1 restricts Salmonella growth through magnesium deprivation,” Science, vol. 366, no. 6468, pp. 995–999, Nov. 2019, doi: 10.1126/science.aax7898.
[5]F. R. Cianfanelli, O. Cunrath, and D. Bumann, “Efficient dual-negative selection for bacterial genome editing,” BMC Microbiol, vol. 20, no. 1, p. 129, May 2020, doi: 10.1186/s12866-020-01819-2.
[6]B. Roche et al., “Heterogeneous dual-metal control of Salmonella infection,” Oct. 24, 2023, bioRxiv. doi: 10.1101/2023.10.23.562652.

#### Supplementary table S4: reagents used in this study.

| Chemical | Abbreviation | MW | Solubilizing agent | Stock solution concentration | Manufacteur | Reference |
|---|---|---|---|---|---|---|
| Iron(III) chloride | $FeCl_3$ | 162.21 | HCl 0.1X | 2.2M | *Prolab* | 24 207.291 |
| Iron-55(III) chloride | $^{55}FeCl_3$ | 162.21 | HCl 0.5X | 2.7mM | *Perkin Elmer* | NEZ043002MC |
| Ferrioxamine E | DFO-E | 653.53 | DMSO | 10mM | *Sigma-Aldrich* | 38266 |
| Ferrioxamine B | DFO-B | 656.79 | DMSO | 10mM | *Sigma-Aldrich* | D9533 |
| Ferrichrome | FC | 687.70 | DMSO | 10mM | *Sigma-Aldrich* | F8014 |
| Enterobactin | ENT | 669.55 | DMSO | 10mM | *Sigma-Aldrich* | E3910 |
| 2,3-Dihydroxybenzoic acid | DHBS | 154.12 | DMSO | 10mM | *Sigma-Aldrich* | 126209 |
| Streptomycin | Strep | 581.57 | H2O | 90mg/ml | *Euromedex* | 1121-A |
| Kanamycin | Kan | 484.5 | H2O | 50mg/ml | *Euromedex* | EU0420 |
| 2,6-Diaminopimelic acid | DAP | 190.20 | DMSO | 10mM | *Sigma-Aldrich* | D1377 |
| Phosphate-buffered saline | PBS | | H2O | 10X | *Sigma-Aldrich* | P7059 |
| Carbonyl cyanide 3-chlorophenylhydrazone | CCCP | 204.62 | DMSO | 100mM | *Sigma-Aldrich* | C2759 |
| Hydrochloric acid | HCl | 36.46 | | 37% | *Sigma-Aldrich* | 258148 |
| Dimethyl sulfoxide | DMSO | 78.13 | | 1X | *Sigma-Aldrich* | 472301 |
| Lysogeny Broth | LB | | H2O | 1X | *Euromedex* | BI-SD7006-250G |
| Ammonium chloride | NH4Cl | 132.14 | H2O | 1X | *Merck* | 101217 |
| Potassium Sulfate | K2SO4 | 174.27 | H2O | 1X | *Carlo Erba* | 363608 |
| Potassium dihydrogen phosphate | KH2PO4 | 136.09 | H2O | 1X | *Carlo Erba* | 361507 |
| Magnesium chloride | MgCl2 | 203.31 | H2O | 1X | *Carlo Erba* | 349377 |
| 4-(2-hydroxyethyl)-1-piperazineethanesulfonic acid | HEPES | 238.30 | H2O | 100mM | *Euromedex* | 10-110-C |
| 2-Morpholinoethanesulphonic acid | MES | 195.24 | H2O | 100mM | *Euromedex* | EU0033 |
